# An improved genome assembly uncovers prolific tandem repeats in Atlantic cod

**DOI:** 10.1101/060921

**Authors:** Ole K. Tørresen, Bastiaan Star, Sissel Jentoft, William B. Reinar, Harald Grove, Jason R. Miller, Brian P. Walenz, James Knight, Jenny M. Ekholm, Paul Peluso, Rolf B. Edvardsen, Ave Tooming-Klunderud, Morten Skage, Sigbjørn Lien, Kjetill S. Jakobsen, Alexander J. Nederbragt

**Affiliations:** Centre for Ecological and Evolutionary Synthesis, Department of Biosciences, University of Oslo., Oslo, Norway.; Department of Natural Sciences, University of Agder, Kristiansand, Norway.; Centre for Integrative Genetics (CIGENE), Department of Animal and Aquacultural Sciences, Norwegian University of Life Sciences, NO-1432, Ås, Norway.; J. Craig Venter Institute, 9704 Medical Center Drive, 20850, Rockville, MD, USA.; Genome Informatics Section, Computational and Statistical Genomics Branch, National Human Genome Research Institute, National Institutes of Health, Bethesda, MD, USA.; Yale School of Medicine, Yale University, 06520, New Haven, CT, USA.; Pacific Biosciences, Menlo Park, CA, USA.; Institute of Marine Research, P.O. Box 1870, Nordnes, NO-5817, Bergen, Norway.; Biomedical Informatics Research Group, Department of informatics, University of Oslo., Oslo, Norway.

**Keywords:** assembly algorithms, assembly consolidation, dinucleotide repeats, *Gadus morhua*, heterozygosity, indel polymorphism, long-read sequencing technology, microsatellites, PacBio, repetitive DNA

## Abstract

**Background:** The first Atlantic cod (*Gadus morhua)* genome assembly published in 2011 was one of the early genome assemblies exclusively based on high-throughput 454 pyrosequencing. Since then, rapid advances in sequencing technologies have led to a multitude of assemblies generated for complex genomes, although many of these are of a fragmented nature with a significant fraction of bases in gaps. The development of long-read sequencing and improved software now enable the generation of more contiguous genome assemblies.

**Results:** By combining data from Illumina, 454 and the longer PacBio sequencing technologies, as well as integrating the results of multiple assembly programs, we have created a substantially improved version of the Atlantic cod genome assembly. The sequence contiguity of this assembly is increased fifty-fold and the proportion of gap-bases has been reduced fifteen-fold. Compared to other vertebrates, the assembly contains an unusual high density of tandem repeats (TRs). Indeed, retrospective analyses reveal that gaps in the first genome assembly were largely associated with these TRs. We show that 21 % of the TRs across the assembly, 19 % in the promoter regions and 12 % in the coding sequences are heterozygous in the sequenced individual.

**Conclusions:** The inclusion of PacBio reads combined with the use of multiple assembly programs drastically improved the Atlantic cod genome assembly by successfully resolving long TRs. The high frequency of heterozygous TRs within or in the vicinity of genes in the genome indicate a considerable standing genomic variation in Atlantic cod populations, which is likely of evolutionary importance.

## Background

The speed and affordability of sequencing and improved software, including more efficient genome assemblers, have led to a democratization of genomics, enabling individual research groups to create *de novo* genome assemblies [1]. The first published *de novo* assemblies for non-model organisms using pure massively parallel sequencing approaches (Illumina and 454) appeared in 2010–2011 and included diverse species such as giant panda [2], turkey [3], woodland strawberry [4] and Atlantic cod [5]. Numerous genome assemblies from a myriad of non-model plants, invertebrates and vertebrates are now available, including examples of genomes that are difficult to assemble, e.g. the extremely large genomes of bread wheat [6] and Norway spruce [7], the highly heterozygous genome of oyster [8] and the tetraploid and repetitive salmon genome [9]. These genome assemblies have provided exciting new biological findings, including the first example of a vertebrate immune system, that of Atlantic cod, which lacks MHC (major histocompatibility complex) class II [5], untangling of the events of multiple hybridizations shaping the ancestral genomes of bread wheat prior to domestication [10] and multiple genomes resolving the avian phylogeny, their radiations and investigation of the genetic basis of complex traits [11, 12]. Despite the tremendous impact of the high throughput sequencing generated genomes, many of these assemblies are of varying completeness, depending on the purpose for which they have been obtained [13, 13]. In the examples given above, the sizes of the scaffold sequences are usually far shorter than chromosome arm lengths. Most of these genomes have scaffold N50 lengths (i.e., half the assembly is in scaffolds of this length or longer) in the range of 400 kbp – 1.5 Mbp, although some avian genomes have N50 scaffold lengths up to 10 Mbp, approaching chromosome arm lengths. However, contig N50 lengths are far shorter and in the range of 3 kbp – 55 kbp.

The presence of repetitive DNA is the most important factor contributing to fragmented genome assemblies [14, 15]. Assembly algorithms might not resolve repetitive regions if they are longer than the read length, and this in particular affects the assembly of sequencing data from short-read technologies such as the Illumina platform [14–16]. Repetitive regions can be divided into two classes, interspersed and tandem repeats. Interspersed repeats, including transposable elements (TEs), occur across the genome and are present in all vertebrate genomes, comprising from 5 % to 55 % of their assemblies [17]. Tandem repeats (TRs) are sequences with a repeat unit repeated more than two times in tandem. Eukaryotic genomes typically consist of 0.5 to 3 % TRs, and TRs can be classified into microsatellites, also called simple repeats, or short tandem repeats (STRs, 1–9 bp repeat unit size); minisatellites (10–100 bp) and satellite repeats (>100 bp repeat unit size) [18]. TRs mutate by adding or removing full repeat units and their mutation rates can be from 10 to 10,000 fold higher than for the remainder of the genome [19]. The heterozygosity caused by TR mutations, in addition to other types of heterozygosity, are also likely to have confounding effects on the contiguity of genome assemblies [14, 15].

Long-read sequencing technologies such as PacBio and Oxford Nanopore address the drawbacks of short-read technologies by enabling read-through of larger repeat regions, and are therefore particularly well-suited for *de novo* genome assembly [14, 20]. Combining moderate amounts of PacBio coverage (5–20x) with other sequencing data can dramatically improve the contiguity of an assembly [21, 21]. More extensive coverage in long reads (>50x) has enabled assemblies of vertebrate genomes to approach complete chromosome arms [23–25], although the associated costs are substantial. A sequencing strategy including long-reads is recommended to aid in reducing the fragmentation typical of *de novo* genome assemblies based on a short-read technology only. Regardless of sequencing strategy, use of a genetic linkage map, or an optical map, can place contigs or scaffolds into chromosome-sized reconstructions, called linkage groups, a prerequisite for large-scale genome comparisons between species [26].

The first release of the Atlantic cod (*Gadus morhua*) genome was sequenced and assembled solely with the 454 sequencing technology [5] and annotated by the Ensembl Project [27] (gadMor1). The 832 Mbp assembly was fragmented, with a contig N50 of 2.3 kbp and 27 % of bases in gaps. The genome assembly contained 17.8 % TEs and 5.9 % TRs (Supplementary Table 6 in [5]). An increased abundance of short unit size TRs at the contig termini (32 %), and at the gaps in scaffolds (24 %, Supplementary Note 7 in [5]) indicate that these repeats contributed to the observed level of fragmentation.

A more contiguous reference genome for Atlantic cod, preferably with chromosome-level reconstructions, will facilitate re-sequencing efforts addressing population genomics investigations, including the detection of structural variants, introgression and hybridization between species, as well as improve comparative genomic investigations relying on synteny. Moreover, it will also enable an annotation with more complete gene models and allow for a better understanding of the lack of sequence contiguity in gadMor1. To achieve this, we created several assemblies using different combinations of Illumina, 454 and PacBio sequencing technologies, Sanger BAC-end sequences, and a suite of assembly programs. As often is the case [28–30], no single assembly outperformed the others in all criteria (N50 contig/scaffold length, gene content, agreement with a genetic linkage map, accordance with read data), thus a reconciled assembly was created to integrate the best characteristics of four draft assemblies. This new assembly (gadMor2) has a fifty-fold improvement of the contig N50 length of gadMor1, and eight times longer scaffold N50 and one sixteenth the number of bases in gaps than gadMor1. A linkage map (personal communication, Sigbj0rn Lien) was used to order and orient the scaffolds into linkage groups. The new genome assembly and annotation reveal a high content of TRs compared to other vertebrates and most notably in promoter regions and amino acid coding sequences. Many of these TRs are heterozygous, and we propose this has implications for understanding local adaptation at a population level.

## Results

An improved genome assembly for Atlantic cod

In addition to already existing sequencing data for the wild-caught individual from the North East-Arctic population described in [5] (˜40x Roche/454 and ˜0.1x Sanger BAC-ends), we added sequencing data from Illumina (˜480x coverage) and PacBio (˜19x coverage) (Supplementary Table 1) obtained from DNA isolated from the same individual. Different assembly strategies were used: a Newbler assembly with 454 and Sanger BAC-end sequences as input (NEWB454), an ALLPATHS-LG [31] assembly with the Illumina sequences only (ALPILM), a Celera Assembler [32] assembly with 454 and Illumina sequences (CA454ILM) and a Celera Assembler assembly with 454 paired reads, Illumina reads and raw, uncorrected PacBio reads (CA454PB) (Supplementary Table 1). For each of the individual assemblies, different combinations of the assembly improvement programs Pilon [33] and PBJelly [34] were applied to improve the consensus sequence and to close gaps (Supplementary Table 2). The properties of these assemblies were assessed using multiple tools: 1) Methods based on the mapping of read datasets to an assembly, *FRC*^*bam*^ [35] and REAPR [36]; 2) by comparing a transcriptome to an assembly, Isoblat (using the Newbler transcriptome, see Methods) [37]; 3) by comparing the assembly to a linkage map (see Methods); 4) and determining presence and completeness of conserved eukaryotic and Actinopterygii (ray-finned fishes) gene sets, CEGMA [38] and BUSCO [39] (Supplementary Table 2).

Based on these evaluations, each assembly had distinct properties, and none is superior for all metrics. For instance, the NEWB454 assembly has the longest scaffold N50 and the lowest number of conflict sequences (Figure 1, Table 1). In contrast, the CA454PB outperforms the other assemblies based on contig N50, yet has a lower scaffold N50 and higher number of sequences conflicting with the linkage map (sequences that map to two linkage groups) (Table 1). Existing assembly reconciliation tools are limited to combining two assemblies [40, 41] and do not perform satisfactorily. To obtain the best possible assembly, i.e., to integrate the information recovered by the different assemblies, we developed a novel assembly reconciliation method. This method involved an all-against-all alignment of the assemblies using Mugsy [42] after splitting the different assemblies in locations where they were in conflict with the linkage map (see Methods) and removing sequences shorter than 1000 bp. The resulting alignment graph structure was traversed following the path from one of the original assemblies (CA454ILM, the one with the most genes found with CEGMA and BUSCO), yielding the sequence from the assembly with the least gaps (CA454PB), while using the alignments with ALPILM and NEWB454 in the graph to close gaps and extend scaffolds. The scaffold module from SGA [43] was applied on the resulting merged assembly using all paired reads (Illumina, 454 and sequenced BAC-ends), and Pilon [33] was used to improve per-base accuracy and to close or reduce gaps. The resulting assembly was ordered and oriented based on a linkage map of 9355 SNPs (personal communication, Sigbj0rn Lien) placing 93 % of the sequences into 23 linkage groups (Supplementary Table 3). Comparisons of assembly statistics for the final, reconciled assembly (gadMor2) and the original four (CA454ILM, CA454PB, ALPILM and NEWB454), show that gadMor2 out-performs all other assemblies on all quality metrics apart from scaffold N50 (ranked 2^nd^) and CEGMA gene content (ranked 3^rd^, Table 1, Figure 1). Based on an overall assessment of quality, gadMor2 combines the best features of each of the four original assemblies without loss of quality (Table 1).

**Figure 1.**
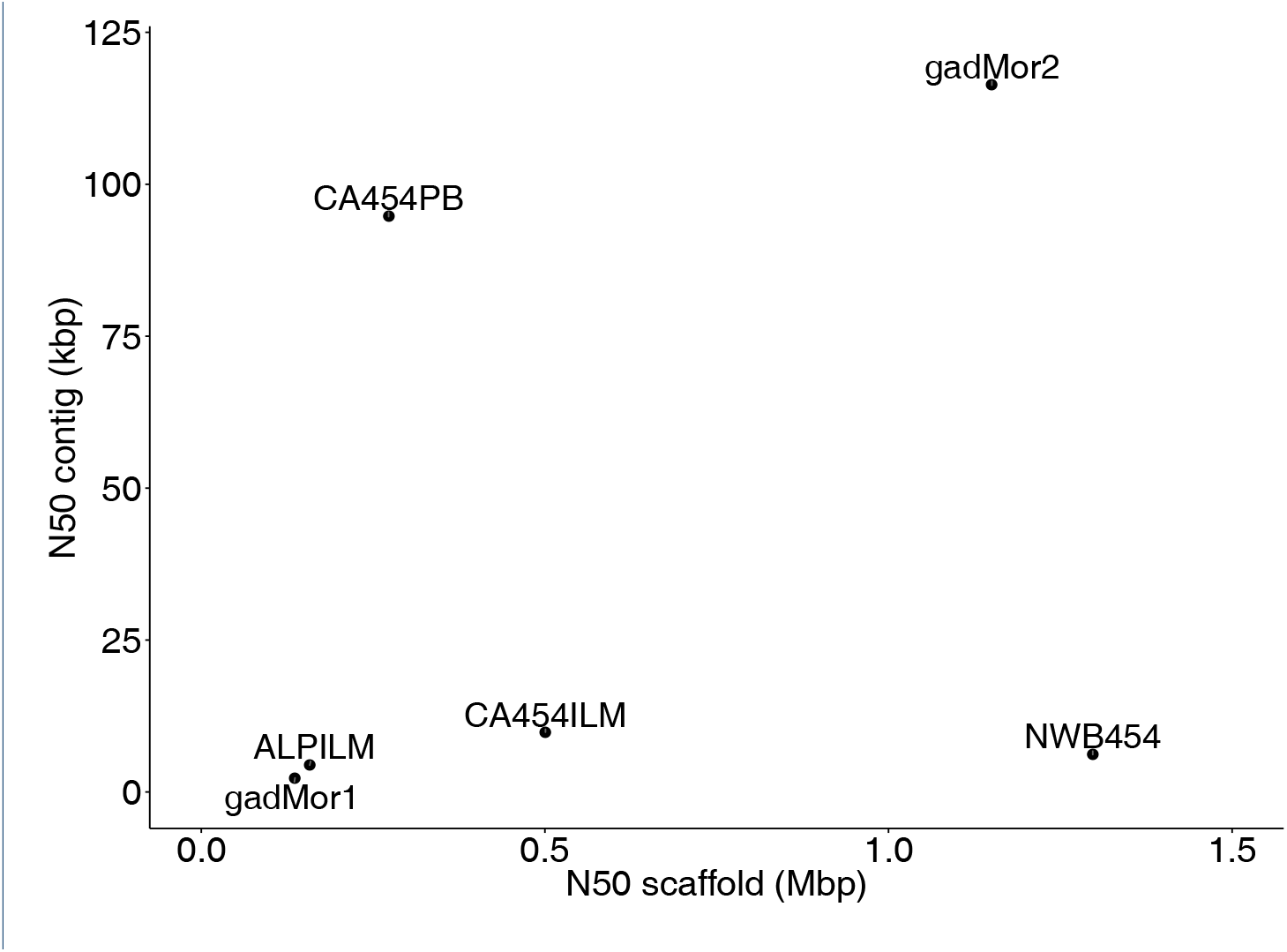
**Contig and scaffold N50 lengths of the different cod assemblies.** gadMor2 was created by following the sequences in CA454ILM in a path through a graph created from a multiple alignment of the four original assemblies, and outputting the contig sequences from CA454PB for each alignment. NEWB454 and ALPILM were used to extend the scaffolds, see Table 1.

**Table 1.** **Overview of assembly statistics.**

| Assembly | Total size assembly (Mbp) | N50 contig (kbp) | N50 scaffold (Mbp) | Percentage gap bases | CEGMA<sup>1</sup> | BUSCO<sup>2</sup> | REAPR<sup>3</sup> | *FRC<sup>bam</sup>*<sup>4</sup> | Potential conflict (sequences)<sup>5</sup> |
| --- | --- | --- | --- | --- | --- | --- | --- | --- | --- |
| gadMor1<sup>6</sup> | 832 | 2.3 | 0.14 | 26.9 | 444 (96.9 %) | 3 308 (89.4 %) | 2 547 | 4 210 772 | 76 |
| ALPILM | 660 | 4.4 | 0.16 | 28.7 | 424 (92.6 %) | 3 016 (81.6 %) | 19 787 | 2 182 096 | 122 |
| NEWB454 | 656 | 6.2 | 1.30 | 24.4 | 435 (95.0 %) | 3 109 (84.1 %) | 18 117 | 2 044 008 | 26 |
| CA454ILM | 647 | 9.9 | 0.50 | 3.49 | 447 (97.5 %) | 3 379 (91.4 %) | 7 406 | 1 351 500 | 96 |
| CA454PB | 682 | 95 | 0.27 | 1.62 | 431 (94.1 %) | 3 310 (89.5 %) | 8 617 | 1 508 054 | 188 |
| gadMor2<sup>7</sup> | 643 | 116 | 1.15 | 1.69 | 435 (95.0 %) | 3 447 (93.2 %) | 7 359 | 1 248 792 | 15 |

<sup>1</sup> CEGMA annotates 458 highly conserved eukaryotic genes
<sup>2</sup> BUSCO annotates 3,698 actinopterygii specific genes
<sup>3</sup> REAPR analyses the discordance between the expected order, orientation and distance of mapped paired reads, with detected potential errors, fewer is better
<sup>4</sup> *FRC<sup>bam</sup>* uses a similar approach as REAPR, with total number of features (i.e., potential assembly problems), fewer is better
<sup>5</sup> number of sequences mapping to more than one linkage group or to multiple linkage groups, fewer is better
<sup>6</sup> from [5]
<sup>7</sup> 93 % of the gadMor2 assembly is additionally oriented and ordered into 23 linkage groups (Supplementary Table 3)

The gadMor2 assembly has a fifty-fold longer contig N50 and eight-fold longer scaffold N50 compared to the gadMor1 assembly [5]. This has dramatic consequences for the sequence contiguity; for instance, a 100kbp region containing the HoxC cluster is a single contig in gadMor2, while it previously consisted of 21 contigs and 20 gaps in gadMor1 (Figure 2).

**Figure 2.**
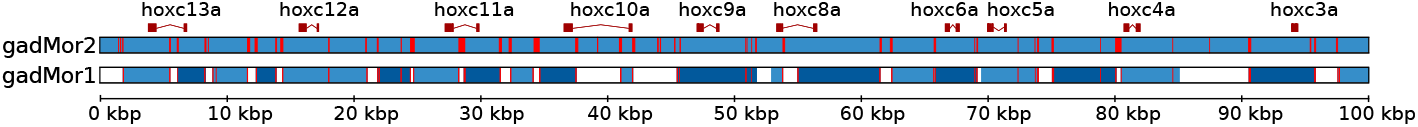
**The HoxC cluster in gadMor1 and gadMor2.** Blocks of dark and light blue are contig sequences, white blocks are gaps and red lines are tandem repeats. Gene models are sketched at the top of the figure. This region is a single contig in gadMor2 and 21 contigs in gadMor1. TRs are at the borders between almost all gaps and contigs in gadMor1.

### Genome size

Estimation of genome size with odd-sized k-mers from 17 to 31 with SGA PreQC [44] on the 300 bp insert size, 100 bp length, paired end Illumina reads (about 150x coverage), resulted in a genome estimate of 613 Mbp±11 Mbp (Supplementary Table 4). The assembler ALLPATHS-LG estimated the genome to be 651 Mbp based on the k-mer distribution of the 180 bp insert size, 100 bp length, paired end Illumina reads (about 52x coverage). Both estimates are lower than previous ones based on Feulgen Image Analysis Densitometry at 0.93 pg or 910 Mbp [45, 45] and a k-mer analysis based on 454 reads, which resulted in a 830 Mbp estimate [5]. Although the assembly size of the gadMor1 at Ensembl is 832 Mbp with 26.9 % gaps [5], the amount of sequence in contigs is 608 Mbp (224 Mbp in gaps), considerably closer to the SGA PreQC estimate. The likely explanation for the large size of gadMor1 is that many of the contigs could not be placed into a scaffold, and a gap was created at that locus instead. These unplaced contigs are included in the output, resulting in loci represented twice in the assembly, once as a gap and once as a contig. The assemblies created in this study all span approximately 650 Mbp, which is similar to the ALLPATHS-LG estimation. 650 Mbp is 71 % of the earlier estimation based on Feulgen Image Analysis Densitometry, and in line with similar results in platyfish (70–89 % of earlier estimations) [47] and in northern pike (64–100 %) [48]. In addition, CEGMA [38] and BUSCO [39] find 89–93 % of conserved genes (Table 1), likely reflecting the approximate completeness of the genome assembly.

### Annotation

We annotated 83,505 gene models with MAKER2 [49, 50], obtaining a final set of 23,243 predicted genes after discarding gene models with low support (see Methods). Compared to gadMor1 (20,095 predictions) [5], the gadMor2 annotation contains more predicted genes and significantly more sequence in the predicted transcriptome (32.2 Mbp and 52.9 Mbp, respectively). The predicted transcripts are substantially longer and without gaps (Table 2). A genome browser enabling access to the genome and the annotation is available [51].

**Table 2.** **Comparison between the gene annotations of gadMor1 and gadMor2.**

| Assembly | Total size transcriptome (Mbp)<sup>1</sup> | Number of genes | N50 length (bp)<sup>2</sup> | Amount gap bases (Mbp)<sup>3</sup> | BUSCO <sup>4</sup> |
| --- | --- | --- | --- | --- | --- |
| gadMor1 | 32.2 (24.8) | 22,618<sup>5</sup> | 1,854 (1,398) | 1.7 | 2,947 (79.7 %) |
| gadMor2 | 52.9 (33.4) | 23,246<sup>6</sup> | 3,239 (1,995) | 0 | 2,714 (73.4 %) |

<sup>1</sup> sum of bases in transcripts with UTRs (without UTRs)
<sup>2</sup> half the transcriptome in sequences of this length or longer, with UTRs (without UTRs)
<sup>3</sup> gaps represented as 'N's in annotated transcripts
<sup>4</sup> number (percentage) of conserved actinopterygii genes detected out of a total of 3,698
<sup>5</sup> when excluding pseudogenes, alternative transcripts, etc., the number of protein-coding genes is 20,095
<sup>6</sup> protein-coding genes only

### Heterozygosity

Illumina paired-end reads with 300 bp insert size and 100 bp read length were mapped to the gadMor2 assembly using BWA-MEM [56], and 2,621,997 SNPs (single nucleotide polymorphisms), 90,292 MNPs (multiple nucleotide polymorphisms), 631,063 indels (insertions and deletions) and 169,181 complex regions (composite insertion and substitution events) with quality ≥20 were called using FreeBayes [57]. With 2,621,997 SNPs, this corresponds to a (SNP) heterozygosity rate of 4.07 χ 10^-3^ (one segregating site every 246 bp). The indel rate in Atlantic cod is 0.98 χ 10^-3^ (one indel every 1020 bp on average, Table 3).

**Table 3.** **Comparison of the SNP and indel rates of selected organisms.**

| Species | SNP rate<br>(SNPs/base) | Indel rate<br>(indels/base) | N50 contig<br>(kbp) | N50 scaffold<br>(Mbp) |
| --- | --- | --- | --- | --- |
| Atlantic cod (gadMor2) | $4.07 \times 10^{-3}$ | $0.98 \times 10^{-3}$ | 116 | 1.15 |
| Stickleback<sup>1</sup> | $1.43 \times 10^{-3}$ | NA | 83.2 | 10.8 |
| Miiuy croaker<sup>2</sup> | $2.24 \times 10^{-3}$ | $0.61 \times 10^{-3}$ | 73.3 | 1.15 |
| Atlantic herring<sup>3</sup> | $3.2 \times 10^{-3}$ | NA | 21.3 | 1.84 |
| *Ciona savignyi*<sup>4</sup> | $46 \times 10^{-3}$ | NA | 12 | 0.192 |
| *Ciona savignyi*<sup>5</sup> | $46 \times 10^{-3}$ | NA | 47 | 0.989 |

<sup>1</sup> from [52]
<sup>2</sup> from [53]
<sup>3</sup> from [54]
<sup>4</sup> from [55]
<sup>5</sup> from [55] with haplotype assembly and merging

We also called indels based on PacBio sequencing reads using blasr [58] and PB-Honey [59]. 70,278 indels of size ≥20 bp were found, at a rate of 0.1 χ 10^-3^ indels/base, or one indel ≥20 bp every 10,000 bp on average.

### Repeat content

We created a repeat library using a combination of RepeatModeler [60], LTRharvest [61], LTRdigest [62] and TransposonPSI [63] and known eukaryotic TE sequences from RepBase [64] (see Methods). This library masked 31.3 % of the genome assembly (Table 4), with 22.9 % classified as interspersed repeats (most often TEs) and 8.0 % as TRs (ranging from dinucletide to hexanucleotide repeats, at least 20 bp long), both classifications higher than for gadMor1 (17.8 % and 5.9 % respectively, Supplementary Table 6 in [5]), indicating a more complete genome assembly.

**Table 4.**
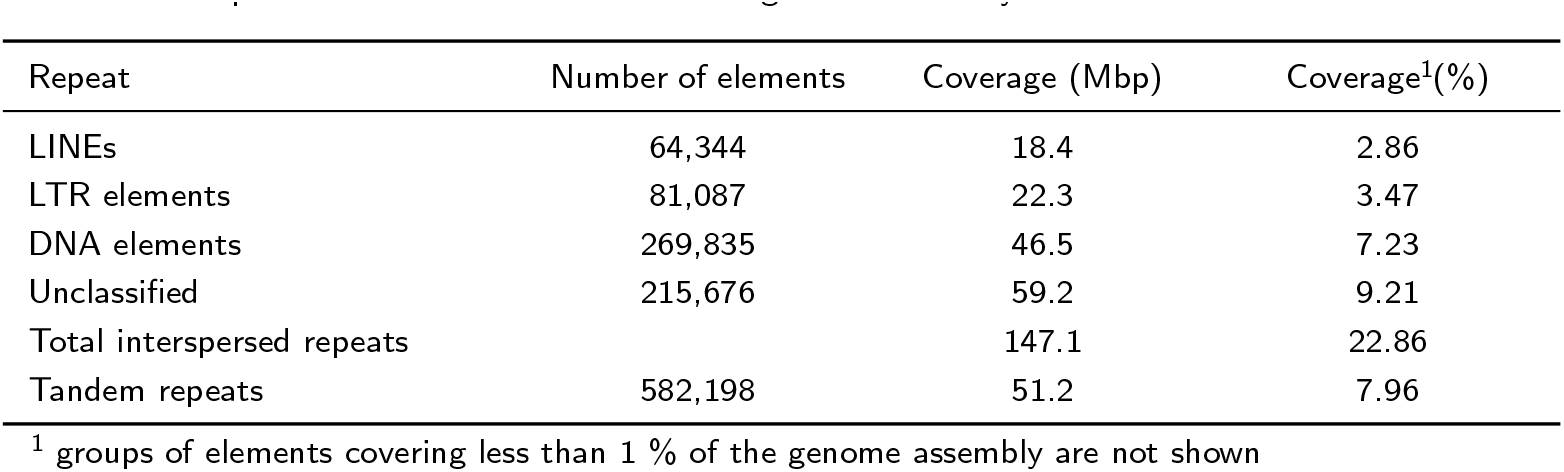
**The repeat content of of the Atlantic cod genome assembly.**

| Repeat | Number of elements | Coverage (Mbp) | Coverage<sup>1</sup>(%) |
| --- | --- | --- | --- |
| LINEs | 64,344 | 18.4 | 2.86 |
| LTR elements | 81,087 | 22.3 | 3.47 |
| DNA elements | 269,835 | 46.5 | 7.23 |
| Unclassified | 215,676 | 59.2 | 9.21 |
| Total interspersed repeats |  | 147.1 | 22.86 |
| Tandem repeats | 582,198 | 51.2 | 7.96 |

<sup>1</sup> groups of elements covering less than 1 % of the genome assembly are not shown

### TR content

We investigated to what extent different assemblers and sequencing technologies affected the numbers of annotated TRs. Phobos [18] was used to find all TRs with a unit size of 1–50 bp, at least 13 bp long (different from the TRs classified above), in the different cod assemblies (Figure 3 and Table 5). Our results shows that assemblies created with the Celera Assembler have the largest amount of TRs (Figure 3).

**Figure 3.**
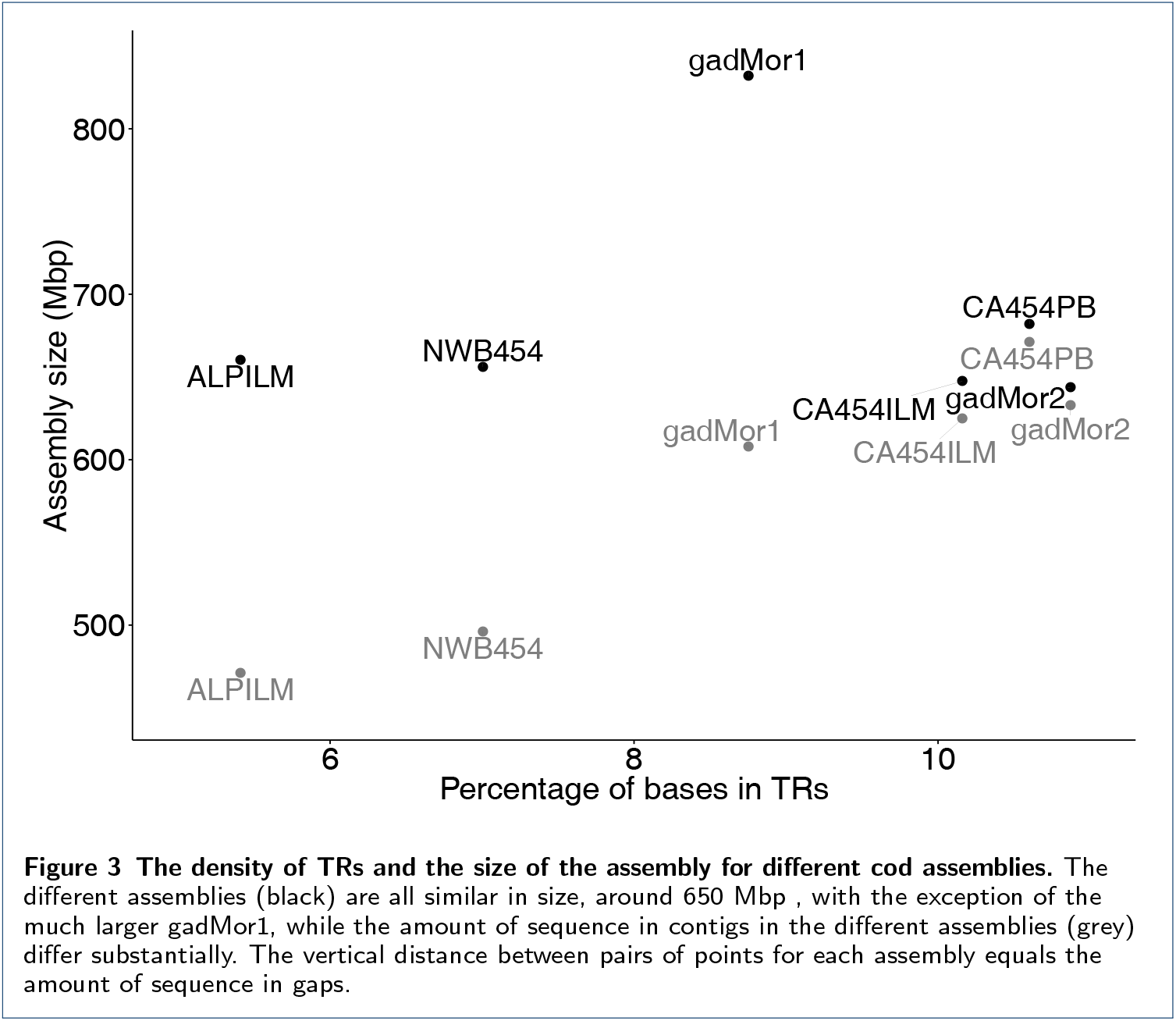
**The density of TRs and the size of the assembly for different cod assemblies.** The different assemblies (black) are all similar in size, around 650 Mbp, with the exception of the much larger gadMor1, while the amount of sequence in contigs in the different assemblies (grey) differ substantially. The vertical distance between pairs of points for each assembly equals the amount of sequence in gaps.

**Table 5.**
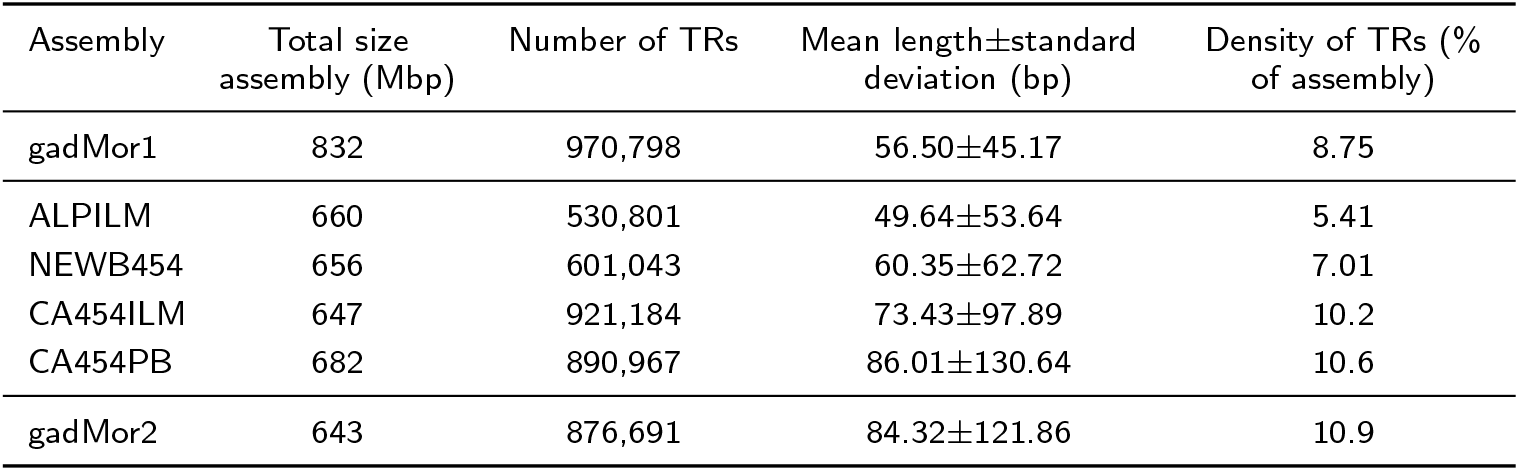
**Overview of tandem repeat statistics.**

The most prominent class of TRs in gadMor2 is dinucleotide TRs, which make up 48.7 % of all annotated repeats, followed by mononucleotide, trinucleotide and tetranucleotide repeats that comprise only 7.6 %, 6.3 % and 6.3 %, respectively (Figure 4). The average length of dinucleotide repeats is 84.4±87.2 bp, at an average 97.3 % identity. In total, dinucleotide repeats make up 5.7 % of the entire gadMor2 assembly. NEWB454 and ALPILM have a significantly lower amount of, and shorter, TRs annotated than the two assemblies created with Celera Assembler, CA454ILM and CA454PB (Table 5).

**Figure 4.**
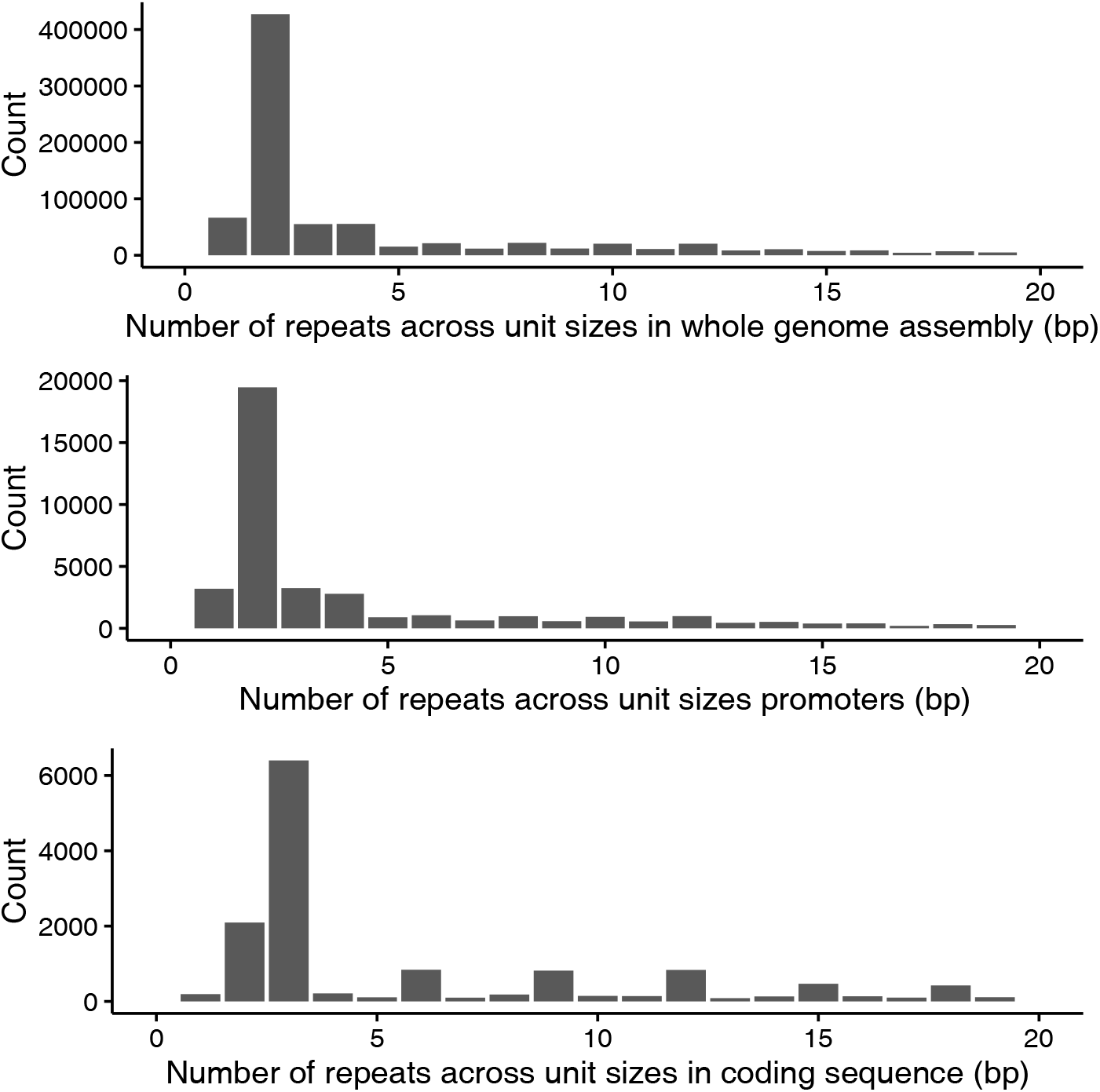
**The number of tandem repeats categorized based on unit size.** Only TRs with unit size 1–20 bp are shown. A unit size of one indicates a mononucleotide tandem repeat, two a dinucleotide, three a trinucleotide, repeats etc. The horizontal axis denotes the unit sizes of the repeat, while the vertical axis shows the count of the particular repeat.

An analysis of gadMor2 compared to all genomes in Ensembl (release 81, ex-cluding gadMorl), including the genome of California sea hare (which contains a large amount of TRs [65]), shows that the Atlantic cod genome assembly has an approximately three-fold higher density of TRs than the genome assemblies of other vertebrates (Figure 5, see also Supplementary Figure 1).

**Figure 5.**
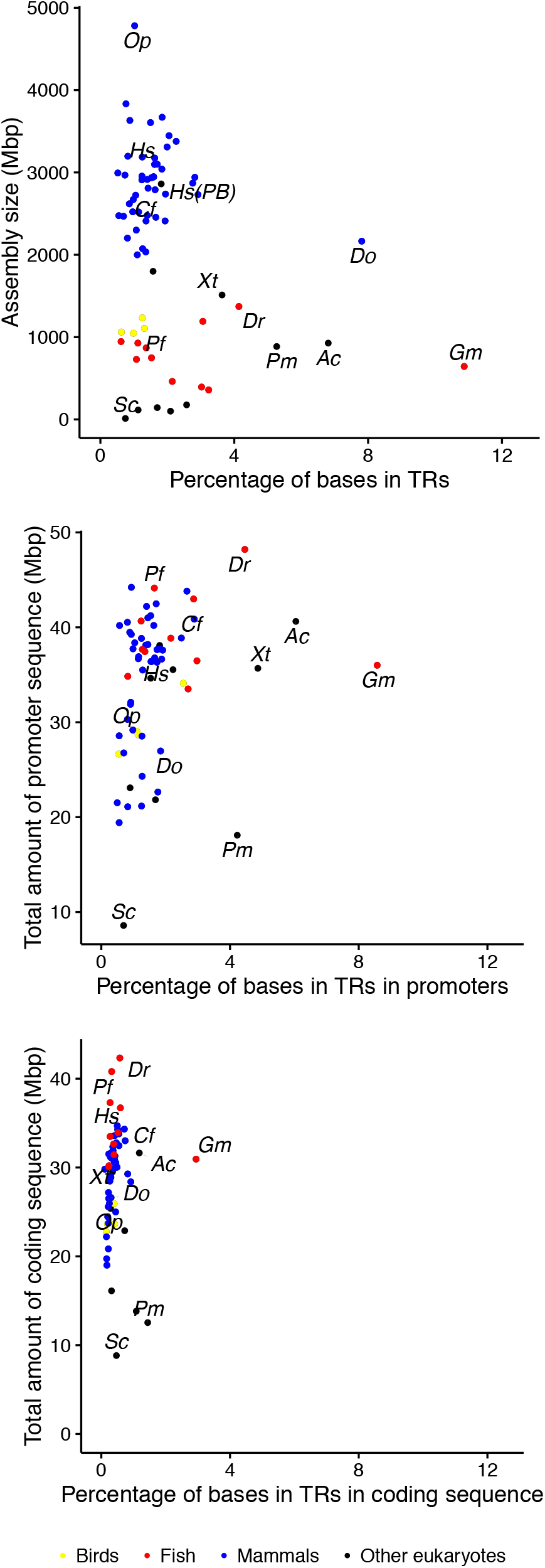
**The density of tandem repeats in genome assemblies, promoters and coding regions.** The assemblies shown here are from Ensembl release 81, excluding gadMor1, plus a human genome based on PacBio data, the California sea hare *Aplysia californica* and gadMor2 (n = 71). The panels show the density (percentage of bases) of TRs in the whole assembly, coding regions and promoter regions, respectively. The human PacBio assembly is not included in the gene and promoter analysis because it has no annotation, and the opossum is lacking for technical limitations. The species marked are *Oc* (*Ochotona princeps*, pika), *Hs* (*Homo sapiens*, human), *Hs* (*PB*) (*Homo sapiens*, human, PacBio based assembly), *Cf* (*Canis familiaris*, dog), *Do* (*Dipodomys ordii*, kangaroo rat), *Xt* (*Xenopus tropicalis*, frog), *Pf* (*Poecilia formosa*, Amazon molly), *Dr* (*Danio rerio*, zebrafish), *Pm* (*Petromyzon marinus*, lamprey), *Sc* (*Saccharomyces cerevisiae*, yeast), *Ac* (*Aplysia californica*, California sea hare) and *Gm* (*Gadus morhua*, Atlantic cod, gadMor2).

### TRs cause fragmentation of non-PacBio based assemblies

To investigate the possible genomic features associated with gaps in APLILM, CA454ILM, CA454PB, NEWB454, gadMorl and gadMor2 assemblies, we mapped the contigs from each assembly to gadMor2 and categorized the intersections between the contig termini (i.e. the positions of the terminal nucleotides of each contig) and different annotated features such as SNPs, indels, TRs, TEs and lack of sequence coverage.

For gadMor2, contig termini overlap most prominently with regions lacking read coverage by any sequencing technology, and annotated TEs. The CA454PB shows the same pattern, albeit with a larger fraction of contig termini not overlapping any annotation, suggesting that these contigs end in large repeats not resolved by any assembly. For the other assemblies, the largest fraction of contig termini overlap with TRs at percentages that are significantly higher (>40 %) than the fraction of the gadMor2 assembly annotated as such repeats (10.9 %, Table 5). As TEs might be longer than the read lengths, they may represent a general challenge for most complex genomes (Figure 6, Supplementary Figure 2).

**Figure 6.**
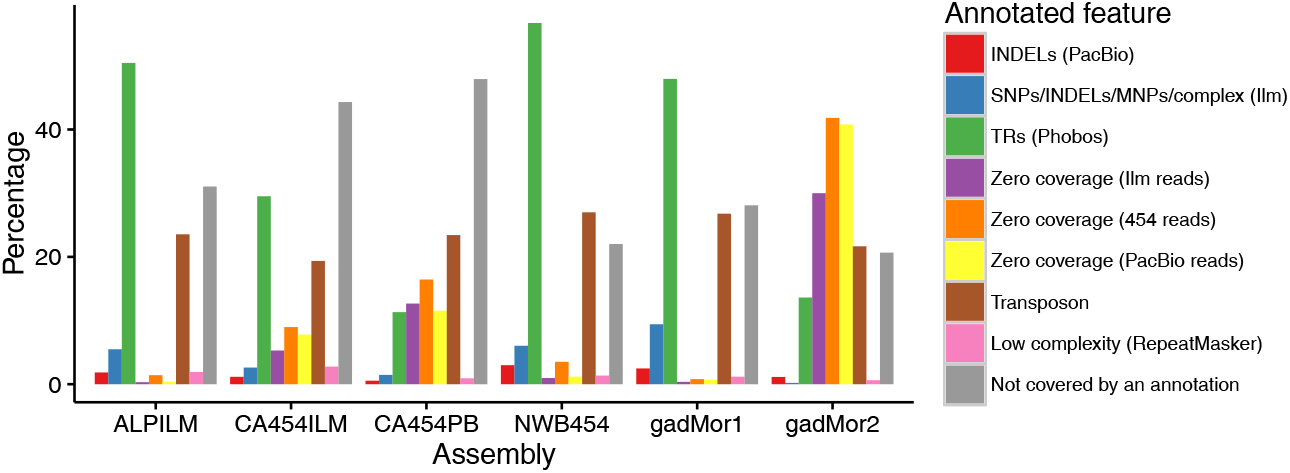
**The intersections between contig termini and different annotated features** The percentage of contig termini (the position of the terminal nucleotides of each contig) intersecting different annotations of the genome.

### Heterozygous TRs

We used lobSTR [66] to investigate the occurrence of heterozygous TRs (i.e., different repeat length between the same locus on the homologous chromosomes) in the sequenced cod genome. lobSTR is designed to analyze TRs with unit length of 1–6 bp (i.e., STRs), and uses Tandem Repeats Finder (TRF) [67] to detect them in the genome assembly. lobSTR both annotates the STRs and discovers variation in STR length. In the sequenced individual, lobSTR annotated 980,400 STRs that passed filtering (1,182,796 in total, see Methods), of which 47,718 were heterozygous.

Compared to Phobos (which annotated 640,938 TRs of units 1–6 bp), lobSTR annotated almost twice as many STRs, and the distributions of the lengths of STRs between the two programs differ largely (Supplementary Figure 3), with lobSTR identifying relatively short STRs, and Phobos annotating relatively long STRs. Given that lobSTR is based on the alignment of the 100 bp Illumina reads, lobSTR's ability to detect heterozygous TRs is limited to repeats around 45 bp in size [68], however, the average length of a TR in cod is 84.32 bp (Table 5). As an alternative to using lobSTR for detecting heterozygous TRs, we used the intersection between TRs annotated by Phobos and indels annotated by either FreeBayes (using Illumina reads, 169,635 intersections) or PBHoney (using mapped PacBio reads, 43,521 intersections). Altogether, 145,435 indels were detected in the 640,938 STRs (1–6 bp unit size) as annotated by Phobos, about three times as many as annotated by lobSTR. For TRs of unit sizes 1–50 bp, there are 183,898 indels in 876,691 TRs (21 %). Our results indicate that at least one-fifth of the TRs in the sequenced individual are heterozygous.

#### TRs in genes and promoters

We investigated the intersection of TRs and coding regions, and found 17,800 coding regions in 7,372 genes contained a TR. Of these TRs, 2,094 TRs (12 %) intersect an indel as annotated by either mapped PacBio or Illumina data. These heterozygous TRs within coding regions are found in 1,514 genes (6.5 % of annotated genes).

In addition, we investigated the 2 kbp sequence upstream of annotated genes (Figure 5). Of the 42,244 TRs identified in these promoter regions, 8,516 (19 %) have an indel annotated based on the union of PacBio and Illumina data.

## Discussion

### An improved genome assembly for Atlantic cod

Here we present a new and significantly improved version of the Atlantic cod genome assembly with successful integration of data from different sequencing technologies. The final assembly (gadMor2) was created using a novel reconciliation method, aimed at combining the strengths of four separate assemblies into an integrated assembly maximizing desired metrics, i.e. contig length, scaffold lengths, gene content and accordance with read data (Table 1). The individual assemblies used for the reconciliation were based on different combinations of sequencing technologies and assembly programs, and varied widely in the different studied metrics. Importantly, the inclusion of the long PacBio reads spanning many more repeats than the other sequencing technologies, resulted in an assembly (CA454PB) with a contig N50 an order of magnitude longer than the other assemblies, contributing directly to the long contig N50 of the final assembly. To our knowledge, the specific approach used in generating CA454PB, where the raw, uncorrected PacBio reads were first trimmed and then assembled without correction, together with Illumina and 454 data (see Methods), has not been previously described. A similar approach was used in generating one assembly for Atlantic salmon (see Supplement in [9]), but the sequence in that assembly did not contribute to the final assembly. End-sequenced BAC (Bacterial Artificial Chromosomes) libraries provide long-range information in the 100 kbp range, and such sequences are available for Atlantic cod [5]. The insert size distribution of the BAC-end library was bi-modal (Supplementary Figure 10 in [5]), which is not handled properly in the Celera Assembler. We therefore included these data in the Newbler assembly (NEWB454) only, which contributed to this assembly having the longest N50 scaffold of the original assemblies. The assembly using a combination of 454 and Illumina sequencing reads (CA454ILM) was the most complete in regards to genes as found by the assembly validation tools CEGMA and BUSCO. While the available Illumina sequencing read datasets did not exactly match the recommendations for ALLPATHS-LG [31], the resulting assembly (ALPILM) performed better than gadMor1 with regards to N50 contig and scaffold metrics. Despite its short contigs and scaffolds, this assembly contributed to the assembly reconciliation process, resulting in longer scaffolds. Our results illustrate a dilemma in obtaining high quality genome assemblies: different combinations of datasets and software using algorithms optimized for certain characteristics of the datasets yield assemblies that are of good quality on different combinations of desired quality criteria, but hardly ever on all [30]. Assembly reconciliation helps solve this issue [41], however even our integrated assembly does not rank best on every single metric evaluated. Further improvements in sequencing technology and assembly algorithms are necessary to resolve this problem in genome assembly.

Due to the fragmented nature of the first version of the Atlantic cod genome, gadMor1, gene-models were reconstructed for the annotation using information from the annotated stickleback genome (i.e., ordering and orienting the contigs based on stickleback gene models), and by manual curation (Supplementary Note 17 in [5]). In contrast, the gadMor2 gene models were automatically annotated directly on the genome assembly. This automated annotation did not annotate pseudogenes, in contrast to the manual curated annotation for gadMor1. The difference in annotation might explain why the CEGMA validation results are slightly lower for the new reference genome, since well-annotated gene models in stickleback would be transferred to gadMor1 (Table 1). The gadMor2 assembly shows fewer indications of potential assembly errors as detected by *FRC*^bam^ and in comparison to the linkage map, but more according to the REAPR program. This difference is associated with longer contigs and scaffolds in gadMor2, which enabled REAPR to estimate more long-range errors. The predicted transcriptome is larger in gadMor2 (Table 2), although more genes are found with BUSCO in the gadMor1 predicted transcriptome. BUSCO is designed to detect genes that are often short (as conserved genes are often short [69]), which means they are more likely put together properly in the gene-model optimized gadMor1 assembly, since longer genes are more likely to be fragmented.

### Causes of fragmentation of cod assemblies

To understand the fragmented nature of gadMor1, we first focused on the rate of heterozygosity, as substantial differences between the homologous chromosomes of diploid organisms can fragment an assembly [55]. We compared the heterozygosity rate of the gadMor2 genome assembly (based on sequencing data obtained from the same individual as gadMor1) to three other fish with genomes for which such data is available, i.e. the miiuy croaker [53], three-spined stickleback [52] and Atlantic herring [54], and to the sea squirt *Ciona savignyi* [55], a species with extremely high heterozygosity (Table 3). The genomes for the fishes have been assembled to high contiguity (Table 3). Although a direct comparison may be confounded by the differences in population structure (in addition to different datasets and programs used [70]), and by a larger uncertainty connected with calling indels correctly than with SNP calls [71], there are substantial differences between the different species. gadMor1 had a N50 contig length of 2.3 kbp (Table 1), substantially shorter than even *Ciona savignyi* which has an order of magnitude higher SNP rate than Atlantic cod. While species with higher SNP rates seem to have shorter N50 contig length (disregarding cod), the sequencing and assembly strategies for the different organisms vary. For gadMor1, the high SNP rate may have had some impact on the fragmentation (Figure 6), but it is not the main explanation.

Different combinations of sequencing technology and assemblers vary in their proportion of TRs present in the resulting genome assembly (Figure 3). Assemblies with higher density in TRs also have more sequence in contigs (i.e., less sequence in gaps), indicating that TRs are more completely assembled. The more fragmented assemblies (ALPILM, NEWB454 and gadMor1) have a lower density of TRs and shorter TRs on average, suggesting that TRs led to fragmentation of the assembly (Table 5). Indeed, these assemblies have a much higher proportion (≥40 %) of contig termini intersecting TRs (Figure 6) than the TR density of 10.9 % in gadMor2 (Table 5). Only CA454PB and (the largely CA454PB derived) gadMor2 have about 10 % of their contig termini intersecting TRs. The remaining gaps in CA454PB and gadMor2 are associated with a lack of sequence coverage and TEs longer than the PacBio read lengths (Figure 6). This illustrates the importance of the availability of the PacBio reads, which was the only read type able to span the multitude of TRs in the genome. As illustrated in Figure 2, gadMor2 has a much higher contiguity, while a large fraction of gaps in gadMor1 are flanked with TRs. Thus, our approach to assemble the genome has addressed the fragmentation affecting the gadMor1 assembly. In conclusion, the high occurrence of TRs in the cod genome has caused the fragmentation of gadMor1 and all assemblies except CA454PB and (the largely CA454PB derived) gadMor2. Without the inclusion of reads obtained from the PacBio technology, or similar sequencing technologies that can span long TRs, assembly of genomes with a high density of TRs, such as the Atlantic cod, to a high sequence contiguity will be significantly more challenging.

### The Atlantic cod genome reveals an extraordinary high density of TRs

We have confirmed and extended previous results showing high genomic densities of STRs in Atlantic cod [65, 72] by comparison with 68 eukaryote genomes (mostly vertebrates, Figure 5). While most of the species studied have fewer than 2.5 % of bases in TRs, California sea hare, kangaroo rat and Atlantic cod have more than 6 % bases in TRs. Atlantic cod has by far the highest density (amount of sequence in TRs) and frequency (the rate of TRs, Supplementary Figure 1) of TRs in the whole genome assembly, coding regions and promoters, with only California sea hare having a higher frequency (but not density) of TRs in promoter regions.

### Potential role of TRs in evolutionary processes in Atlantic cod

The mutation rates of TRs, and especially STRs, are orders of magnitude higher than those of other genomic sequences [19, 73, 74]. In the sequenced individual, we find that one fifth of the annotated TRs are heterozygous, with somewhat lower proportions in promoters (19 %) and coding regions (12 %). These results are based on the mapping of Illumina and PacBio reads, but are likely underestimates. Most of the TRs in cod have a short repeat unit that mutate by adding or removing at least one repeat unit, for instance, two nucleotides in the case of dinucleotide repeats. Small differences between two long alleles of a TR would likely not be captured by our analyses, because the Illumina reads would not map well to these [15, 75], and the PacBio reads might not give sufficient resolution.

In humans, TRs are best known in connection with diseases such as Huntington's Disease [76]. In other species, variability (multiple alleles at a locus within a population) in TRs in promoter regions has been associated with diverse phenomena such as behavior in voles [77] and to skull form in dogs [78]. In both *Saccharomyces cerevisiae* and humans, some promoter regions contain TRs [79, 80], for which variation in length has been linked to variation in expression [80, 81]. TRs in promoter regions may also contribute to expression divergence in great apes [82] and speciation in primates [83]. There is also variability in TRs in genes leading to functional variation such as in *Saccharomyces cerevisiae*, where TRs in cell-wall genes underlie variation that causes alterations in phenotype, with different genotypes have differences in adhesion, flocculation or biofilm formation [84]. Further, in Hawaiian mints, variation in a gene coding for a flowering time protein is associated with colonization and radiation of the plant, with longer versions of the gene existing in younger populations and this is suggested to contribute to morphological change and speciation [85]. Interestingly, Atlantic cod has a higher frequency of TRs than these species within both promoters and coding regions (Supplementary Figure 1).

The sequenced individual was from the North-East Arctic cod population, the largest cod population in the world [86], with a large effective population size [87]. Extrapolating the high mutation rate of TRs, and the observed level of polymorphism in this single individual, suggests that most TRs are polymorphic at a population level. These polymorphic TRs contribute substantially to standing levels of genomic variation in Atlantic cod populations within and in the vicinity of genes.

## Conclusions

Atlantic cod has an extraordinary amount of TRs compared to other species. This repeat content has previously led to complications for assembling the genome. Here, this has been addressed with the usage of the long PacBio sequencing reads and reconciliation of multiple assemblies. The large amount of TRs is likely to have profound evolutionary impact. In particular, the TRs in coding and regulatory regions will drive genetic variation affecting the function or regulation of genes in Atlantic cod populations. It remains to be investigated how cod populations evolve under variable environmental conditions with respect to TRs, and whether selection for repeat variation can lead to rapid evolutionary adaptations.

## Methods

### Sequencing

All read datasets originated from DNA extracted from the same individual fish, designated NEAC_001, a wild-caught male specimen of the North-East Arctic population, sampled with the main purpose for sequencing initiative of the Atlantic cod genome and described in detail in [5]. We always strive to limit the effect of our sampling needs on populations and individuals. This individual was sampled in connection with a research survey conducted by Norwegian Institute for Water Research as part of part of larger hauls for stock assessments. The fish were humanely sacrificed by administration of other sedatives before sampling in accordance with the guidelines set by the 'Norwegian consensus platform for replacement, reduction and refinement of animal experiments' (www.norecopa.no). See Supplementary Table 1 for an overview of different DNA datasets generated from this individual.

Roche/454 reads were sequenced as described previously [5]. The Roche/454 software gsRunProcessor version 2.6 was used to redo basecalling for all sequencing runs generated for the NEAC_001 sample [5].

180 bp insert size and 300 bp insert size libraries were constructed with Illumina DNA paired end sample preparation reagents and sequenced at the Norwegian Sequencing Centre. The 5 kbp insert size libraries were prepared with the Illumina Mate Pair gDNA reagents and sequenced at the McGill University and Genome Quebec Innovation Centre. All Illumina libraries were sequenced on the HiSeq 2000 using V3 chemistry 100 bp paired end reagents.

PacBio SMRT sequencing was performed on PacBio RS instrument (Pacific Biosciences of California Inc., Menlo Park, CA, USA) at the Norwegian Sequencing Centre (www.sequencing.uio.no/) and at Menlo Park. Long insert SMRTbell template libraries were prepared at NSC (10 kbp insert size) and Menlo Park (22 kbp insert size) according to PacBio protocols. In total, 147 SMRT-cells were sequenced using C2 and XL polymerase binding and C2 and XL sequencing kits with 120 min acquisition. Approximately 7.6 Gb of library bases were produced from 10 kb SMRTbell libraries sequenced on 102 SMRT cells using C2/C2 chemistry (average polymerase read length of 3 kb). The 22 kb SMRTbell library was sequenced using C2/XL (22 SMRT cells, average polymerase read length of 4.5 kb) and XL/XL (23 SMRT cells, average polymerase read length of 5 kb) chemistry producing 5.5 Gb of library bases.

### Assembly

An overview of the usage of different sequencing data in the different assemblies is in Supplementary Table 1.

#### ALLPATHS-LG assembly, ALPILM

An ALLPATHS-LG [31] assembly was created using only the Illumina reads. Paired end 100 bp Illumina reads from a 180 bp insert size library were input as fragment reads, while paired end 100 bp reads from a 300 bp insert library and 100 bp reads from a 5k mate pair library were input as jumping reads. Only half of the fragment reads were used in the assembly to have the recommended coverage (Supplementary Table 1). The release R48639 of ALLPATHS-LG was used.

#### Newbler assembly, NEWB454

Newbler version 3.0 was used to assemble the 454 sequencing data together with BAC-ends previously generated for [5], with the options “-large-het-repfill-sio-info-a 0”. In contrast to the Newbler assembly done for the first version of the Atlantic cod genome [5], we did not filter out 454 reads consisting entirely of short TRs, as newer versions of the Newbler program are better able to deal with these reads.

In its output, Newbler gives a file with all scaffolds, including all unscaffolded contigs longer than 2 kbp, and a separate file with all contigs, regardless of their inclusion in a scaffold. Using BLAT version 3.5 [88] we mapped the flanking sequences of SNPs in the linkage map (personal communication, Sigbj0rn Lien) (n=9355) to all contigs. For each mapped SNP, the longest contig to which it mapped was added to the primary output, with the rationale that sequences with SNPs should be included in the assembly. The final assembly thus contains all scaffolds, all contigs longer than 2 kbp and the longest unplaced contigs with a mapped SNP.

#### Celera Assembler assembly based on 454 and Illumina reads, CA454ILM

Celera Assembler's meryl (SVN snapshot dated 2nd of April 2013) [32] was used to count k-mers in the two paired end Illumina read libraries, of 180 bp and 300 bp insert sizes and of length 100 bp.

FLASH version 1.2.3 [89] was used to merge the overlapping reads from the 180 bp library using default options.

The merTrim program, also from Celera Assembler, was used to correct Illumina reads by changing infrequent k-mers to frequent k-mers: starting from the first (last) frequent k-mer in a read, if the next (previous) k-mer is infrequent, then the most recently added base must be an error. To correct it, the three substitution changes are tested; if all k-mers spanning this base are now frequent, the change is accepted. If not, the four insertion and one deletion changes are tested; likewise, if all k-mers spanning this change are now frequent, the change is accepted. Otherwise, the base is left unchanged. Finally, the read is trimmed to the largest region with all k-mers designated as frequent k-mers.

Celera Assembler was used to remove duplicate reads from the 300 bp and 5 kbp Illumina reads libraries with its run runCA-dedupe pipeline.

All 454 reads were converted from .sff files to .fastq and .frg files using Celera Assembler's sffToCA with options “-linker flx-linker titanium-insertsize ins_size std_ins_size-trim chop-libraryname lib_name-output output_name”, with insert sizes and standard deviations at 1100, 320; 1230, 350; 1440, 440; 1760, 470; 2650, 700; 7000, 1900; 19000, 4750 for the different sequencing libraries increasing in insert size (Supplementary Table 1). The insert sizes and standard deviations were those reported by Newbler.

The 454 reads were error-corrected using the merTim program, as above, and trimmed as described in Prüfer et al. [90], removing duplicated pairs of reads, error-prone ends of reads, reads with sequence not confirmed by other reads and chimeric reads. Because the insert length distribution of the paired reads from the 20 kbp 454 mate pair library showed a bimodal distribution (Supplementary Figure 4, in [5]), and since Illumina mate pair libraries contain contamination with pair of reads with the opposite orientation, the scaffolds from this assembly were used to filter out reads from the 20 kbp 454 library and the 5k Illumina library by mapping the reads to the scaffolds using BWA-MEM [56], and removing any pair of reads that mapped closer than 10 kbp and 2 kbp, respectively.

After the error correction steps, all 5 kbp mate pair Illumina reads, 6x coverage of the 300 bp insert size Illumina reads and 25x of the merged 180 bp insert size Illumina reads were assembled together with all the 454 reads. Seqtk [91] from November 2012 was used to extract these reads.

The assembly used this spec file (only non-default options shown):

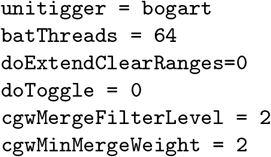

Contigs from Celera Assembler's degenerate contig file, normally excluded from scaffolds, were added to the assembly if they contained flanking sequence from a SNP from the SNP-chip as described above for the Newbler assembly.

#### Celera Assembler assembly based on PacBio, 454 and Illumina reads, CA454PB

All processing of Illumina and 454 reads were redone as described above, using Celera Assembler 8.1.

Filtered subreads of PacBio reads were trimmed using Celera Assembler 8.2 alpha with this spec file (only non-default options shown):

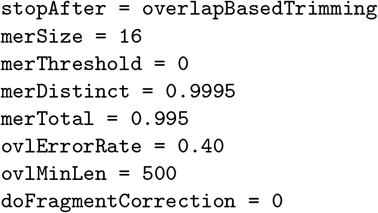

Assembly below was run with this spec file (only non-default options shown):

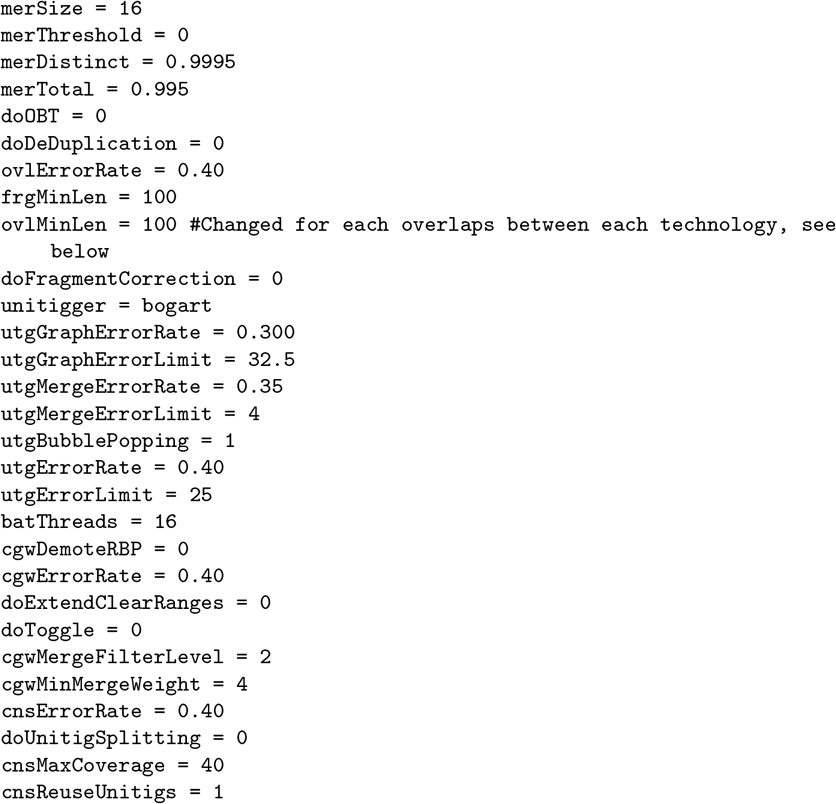

The assembly contains all paired 454 reads, 25x of merged reads from the 180 bp insert size Illumina library and the trimmed PacBio reads, and was run with Celera Assembler 8.2 alpha. To accommodate vastly different error rates between the Illumina/454 and PacBio reads, overlaps were computed using a different percentage maximum allowed error (inverse of percentage identity) cutoff for each pair of technologies being overlapped. Overlaps between Illumina and 454 reads were computed to a maximum of 6 % error and minimum overlap of 100 bp; overlaps between an Illumina/454 read and a PacBio read was computed to a maximum of 20 % error, also with a minimum overlap of 100 bp; overlaps between two PacBio reads were computed to a maximum of 40 % error and minimum overlap of 1000 bp. For each read end, the bogart unitig construction algorithm will pick the longest overlap and use only those for constructing initial unitigs, similar to the BOG algorithm in [32]. Bogart uses clusters of partially aligned reads (discovered via pre-computed overlaps) to detect junctions between repeat and non-repeat sequence. If a detected repeat is spanned by either a read or a mate-pair, the repeat is left intact, otherwise, the unitig is split into at least three pieces: one for each side of the repeat, and at least one for the repeat itself.

The rest of the assembly process was run as normal, aside from much higher error rate acceptance at all steps and a non-default selection of unique unitigs. Because PacBio reads confuse Celera Assembler's classification of unique unitigs (which can be used as seeds for creating contigs) and non-unique unitigs (often repeats that could be placed several times in the assembly), we ran the classification tool markRepeatUnique by hand, specifying that unique unitigs could not have a single reads spanning more than 90 % of its length, up to 15 % of the unitig could have a depth of only 3 reads, and must have had at least 200 reads and be at least 10,000 bp long. Command:

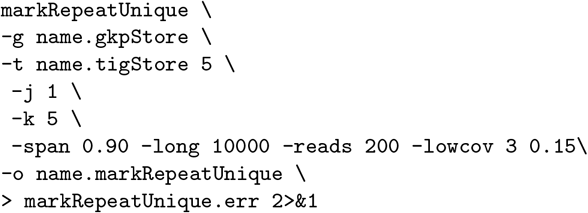

Degenerate sequences that either contained a SNP (as described earlier) or a gene found with CEGMA version 2.4.010312 [38, 69], were added to the assembly output.

#### Pilon and PBJelly

All four assemblies described above were processed with PBJelly (SVN snapshot 23rd September 2014) [34], a tool that maps PacBio reads back to the assembly and uses them to close gaps both between and within scaffolds. The content of the configuration file Protocol.xml:

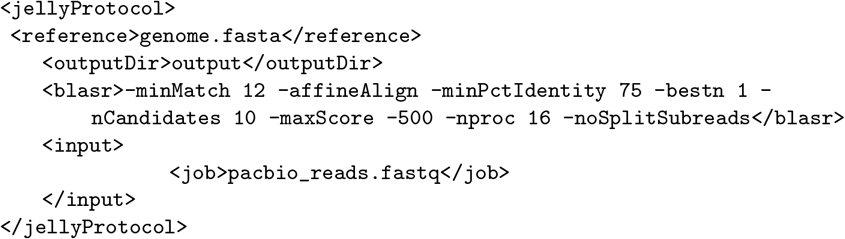

Commands used:

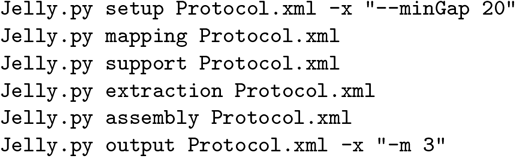

Pilon version 1.9, a program to automatically improve assemblies [33], was applied to both the original and the PBJelly version of the assemblies, using all 454 reads, the reads from the 300 bp and 5 kbp insert size Illumina libraries, mapped with BWA-MEM 0.7.9a and sorted by samtools 0.1.19 [92]:

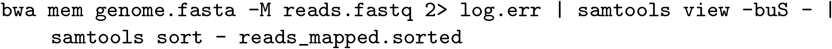

Pilon options were (not showing all the libraries):

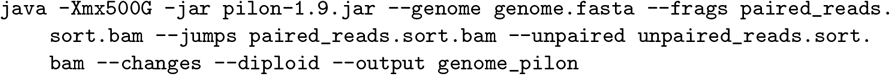

And the reads from all PacBio libraries, mapped with blasr from SMRTanalysis 2.2.0 and sorted by samtools 0.1.19:

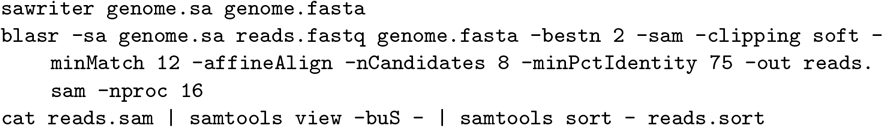

This resulted in four different versions of each assembly: the original; one processed with PBJelly; one processed with Pilon; one and processed with both PBJelly and Pilon. Based on the results of the validation tools against applied to all versions of the assemblies (see below), one version of each assembly was chosen for merging, the versions of ALPILM, NEWB454 and CA454PB after application of both PBJelly and Pilon and the version of CA454ILM after application of Pilon only.

#### Validation

To evaluate assembly quality, several validation tools were applied. Both REAPR [36] and *FRC*^bam^ [35] use paired Illumina reads to evaluate an assembly, giving a measure of the number of potential errors. Instead of using the raw reads, we used error corrected reads dumped from the ALLPATHS-LG assembly, reducing the running time of both the alignment step and the tools themselves.

Isoblat was used to determine how much of the Newbler transcriptome of 454 and Sanger reads was aligned to the different assemblies [37]. It was run with default options.

CEGMA is a tool that annotates 458 highly conserved genes in an assembly, and it can be used to assess the completeness of the genome assembly [38, 69]. Version 2.4 was applied to all different versions of the assemblies.

BUSCO is similar to CEGMA in that it assesses the completeness of a genome by trying to find a set of universal single-copy orthologs [39]. In this study, we used the actinopterygii specific set of 3698 genes to investigate the completeness of the assemblies generated here.

A linkage map for Atlantic cod has been created from a set of 9355 SNPs (personal communication, Sigbj0rn Lien). We used blat_parse.py to compare the linkage map to different assemblies to evaluate the completeness and long-range correctness. Briefly, this involved mapping the flanking sequences of the SNPs to the assembly using BLAT version 3.5 [88] and options “-noHead-maxIntron=100 genome.fasta flanking_sequences.fasta” and then parsing the output file while comparing with the order of the SNPs in the linkage map. A conflict with the linkage map is defined as a sequence that had SNPs mapped to it belonging to more than one linkage group. Some SNPs mapped equally well to more than one linkage group, and these were excluded since we could not confidently judge which mapping was correct.

### Merging of assemblies

Each assembly was aligned against itself using nucmer [93], and any sequences fully contained in another sequence with more than 98 % identity were removed. Scaffolds were split with a split_asm_lg.py (available on the github repository together with the other scripts mentioned in this section) if they conflicted with the linkage map. A scaffold in conflict is split into three pieces, from the start of the scaffold following one linkage group to the last basepair in the flanking sequence of the last SNP in that linkage group, and from the first basepair in the flanking sequence of the first SNP in another linkage group. The middle piece is not used since we do not know where exactly the transition from linkage group to another happens. Sequences shorter than 1000 bp were removed to better facilitate the whole assembly alignment process.

The four assemblies selected for merging were aligned together using Mugsy. Mugsy uses nucmer from the Mummer package [93] to find similar sequence in different assemblies and subsequently refines the alignment. It outputs a MAF (Multiple Alignment Format) file, consisting of blocks of multiple alignments with information where exactly in the sequences the alignment is (starting at 100 bp and ending at 300 bp in scaffold X in assembly Y for instance). The MAF file was parsed by merge_asms.py. Based on the validation criteria described above, one assembly was chosen as the skeleton (CA454ILM), and a second assembly was chosen as the sequence contributing part (CA454PB). The CA454ILM assembly was chosen as skeleton because it was the most complete with regards to genes, and CA454PB was chosen as sequencing contribution assembly was chosen because it had the least gaps. A first pass through the alignment blocks of the first assembly was used to close gaps using the sequences from the CA454PB assembly, or the sequence in each alignment block with the least amount of missing bases. A second pass through the alignment blocks of the first assembly tried to connect scaffolds from the first assembly (CA454ILM) using scaffolds from other assemblies spanning two scaffolds in CA454ILM. Mugsy was run with these options:

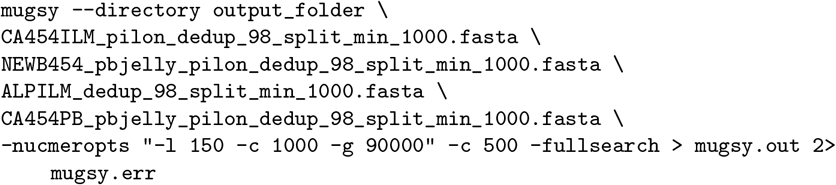

We mapped all paired Illumina and 454 reads to the assembly with BWA-MEM 0.7.9a, and used the scaffold module from SGA [43] to scaffold the merged assembly, increasing N50 scaffold from 850 kbp to 1.15 Mbp. Pilon was then applied using all reads excluding PacBio and the 180 bp insert size Illumina library.

### Anchoring to linkage map

Finally, the scaffolds were ordered into linkage groups based on linkage data (personal communication, Sigbj0rn Lien) with 100 Ns between two adjacent scaffolds using order_orient_scaffolds.py. Scaffolds with only one SNP kept their existing orientation, while scaffolds with more than one SNP were reverse complemented if more than half the SNPs suggested this. The numbering of the linkage groups is according to Hubert et al. [94].

### Transcriptome assemblies

We obtained transcriptome datasets from three different sequencing technologies, Illumina, 454 and PacBio, from a variety of tissues and different stages. Three different transcriptome assemblies were created: (i) based on assembly of the Illumina reads using Trinity [95]; (ii) assembly of the 454 reads using Newbler [96]; and (iii) clustering the long full-isoform PacBio reads using SMRT-Analysis [97].

#### Trinity with Illumina reads

RNA-seq sequencing data used in Penglase et al. [98] (from larvae at different stages and feeding regimes) were obtained from the Sequence Read Archive (SRA) at NCBI with accession ID SRP056073, and adapters and all bases with less than 20 in Phred quality score were removed with cutadapt 1.5 [99]. Trinity version r20140717 [95, 100] was run with the normalize_reads option turned on. 654,948 transcripts were assembled. Abundance estimates commands:

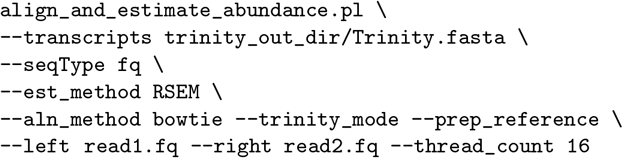

The script filter_fasta_by_rsem_values.pl distributed with Trinity was used to filter the transcript assembly based on abundance, where only transcripts with fragments per kilobase of transcript per million mapped reads (FPKM) of at least 0.05, and a transcript abundance of at least 1 % of the parent gene's abundance were kept, resulting in 59,379 transcripts.

#### Newbler with 454 and Sanger reads

The transcriptome 454 and Sanger reads used in Star et al. [5] (the different tissues listed in Supplementary Table 2 in [5]) were combined with Sanger reads from Kleppe et al. [101], and assembled with Newbler 3.0 using the options -cdna and -vt with these primer sequences:

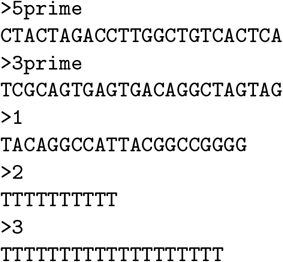

The assembly resulted in 79,025 transcripts.

#### IsoSeq on PacBio reads

Equal amounts of RNA were isolated from pools of unfertilized eggs and at 20, 30, 45, 60 and 90 days post hatch. Isolations were pooled and three size-selected fractions based on agarose gel-electrophoresis of RNA were created at sizes 1–2 kbp, 2–3 kbp and 3–6 kbp and sequenced on the Pacific Biosciences RS using P6v2-C4 chemistry [97]. Using SMRT Portal, reads-of-insert were first created for each fraction, and isoform prediction and polishing by Quiver were performed according to the manufacturer's instructions. For the fraction 1–2 kbp, 10,738 high quality isoforms were predicted (≥99 % accurate sequence according to Quiver) and 2,952 low quality (<99 % accurate sequence), for the 2–3 kbp fraction 15,688 high quality and 6,898 low quality and for the 3–6 kbp fraction 13,400 high quality and 12,716 low quality transcripts. These 62,392 transcripts were merged into one fasta file and used in further analyses.

### Annotation

#### Repeat libraries

A repeat library for MAKER gene annotation (see below) was created by running RepeatModeler [60] version 1.0.8 on the finished genome assembly with default options.

We also created a repeat library specifically for annotation of transposable elements (https://github.com/uio-cels/Repeats). First, RepeatModeler [60] version 1.0.8 was run on only the scaffolds longer than N50. LTRharvest [61] and LTRdigest [62], both parts of genometools (version 1.5.7), were used to detect LTR retrotransposons and TRIMs. LTRharvest found LTR retrotransposons with LTRs larger than 100 nt, smaller than 6000 nt and with 1500 to 25000 nt between, with a target site duplication (TSD) length of 5 nt. TRIMs were detected by lowering the LTR length requirements to a minimum of 70 nt and a maximum of 500 nt with maximum 1500 nt of internal sequence. Harvested putative LTR retrotransposons were filtered using LTRdigest, which checked for tRNA binding sites. In addition, LTRdigest used Hidden Markov Model (HMM) profiles to identify retrotransposon enzymes (from the GyDB HMM profile collection of retrotransposon specific enzymes [102]). Elements without both tRNA binding sites and a retrotransposon specific enzyme were discarded.

We used scripts provided by Ning Jiang, Megan Bowman and Kevin Childs (Michigan State University) to perform the next analyses [49, 103]. Only elements containing primer binding sites (PBS) and/or a polypurine tract (PPT) were kept, and only if at least half of the PBS or PPT sequence was located in the internal regions of the putative element and the distance between the LTRs and the PPT/PBS sequence was less than 20 bp. Elements that passed this filtering were subjugated to further filtering where sequences with gaps of ≥50 nt were discarded. MUSCLE version 3.8.31 [104] was used to align flanking sequences, and elements with ≥60 % similarity in flanking sequences was excluded.

Nested LTR retrotransposons were detected by using RepeatMasker with the left LTR sequences of the putative elements and a library of transposases (from a curated library included in the software TEseeker v1.04 [105]). Consensus sequences were produced after all vs. all comparisons using BLASTN. Finally, no elements from different families shared 80 % sequence over 90 % of their length.

RepeatClassifier, which is a program included with RepeatModeler, was used to classify the elements. As many LTR retrotransposons and TRIMs contain TRs in their long terminal repeats, RepeatClassifier classified some elements as being TRs. These elements were renamed to being LTR retrotransposons or TRIMs, while those that were classified into specific LTR families kept their new classification. TransposonPSI [100] was also run. TransposonPSI uses PSI-BLAST to detect distant homology between genomic sequences and a TE library bundled with the program. Contrary to the other programs, TransposonPSI does not output the consensus sequences of elements detected, which made it necessary to perform an additional clustering step. The output sequences were clustered using CD-HIT-EST 4.6.4 [106] with a similarity cutoff of 80 %. The relative high amount of dinucleotide repeats in the Atlantic cod genome assembly, led to a large fraction of sequence being labeled as transposons of the CACTA superfamily, as the CACTA representative in the TransposonPSI library contained a TR that spurred false alignments. Thus, elements were only named CACTA if two sources agreed in the classification, the other source being the results of a BLASTX search against the repeat peptide database provided with RepeatMasker (version 4.0.6).

As the detection tools might detect repetitive non-TE genes such as gene families, the sequences were checked for alignments (using BLASTX) with sequences in the curated protein database of UniProtKB/SwissProt [107], which was downloaded November 20th 2015. Sequences were also checked against the repeat peptide database that comes with distributions of the RepeatMasker software. Sequences with matches in the UniProtKB/SwissProt database, but not in the repeat peptide database were discarded. The BLASTX search against repeat peptides in the database also served to classify some of the unclassified elements.

Some sequences remained unclassified, and a collection of HMM profiles was downloaded from the Dfam database (Dfam.org) and HMMER3 was run using the nhmmer module. This further classified some elements into LTR retrotransposons, LINEs, SINEs or DNA transposons. The *de novo* library was merged with known eukaryotic repeat sequences from RepBase [64] (version 20150807) and served as input for RepeatMasker.

#### Annotation with MAKER

MAKER is an annotation pipeline designed to combine the consolidated output from different *ab initio* gene finders and physical evidence (e.g. protein and RNA-seq alignments) into a set of quality scored gene models (AED score) [49, 50, 108].

A two-pass iteration with MAKER version 2.31.8 [49, 50] was performed on the final genome assembly as described in [109] and in Campbell et al. [110]. First, two *ab initio* gene finders were trained, SNAP version 20131129 [111] on the genes found by CEGMA version 2.4.010312, and GeneMark-ES version 2.3e [112] on the genome assembly itself. SwissProt/UniProtKB [107] was downloaded 9th of May 2015 (release 2015_04). MAKER was configured to use the two trained *ab initio* gene finders, the SwissProt/UniProtKB protein database [107], the RepeatModeler repeat library and three different transcriptomes, one based on 454 and Sanger data, one based on Illumina and one based on PacBio. Additional options were these:

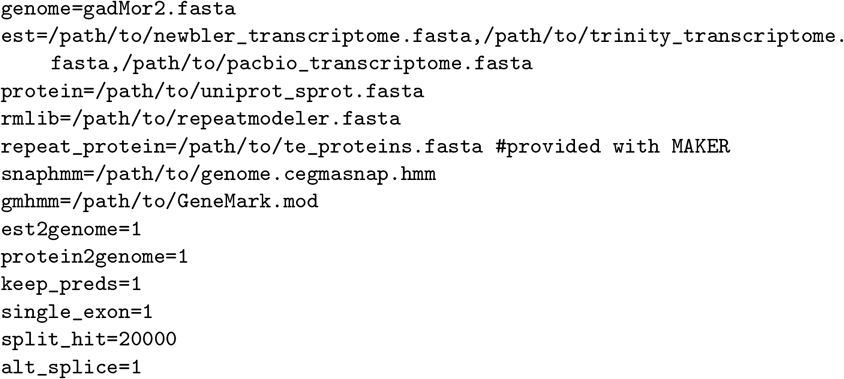

The GFF output from the first pass with MAKER was used to retrain SNAP, and to train AUGUSTUS version 3.0.2 [113, 114] with the PacBio transcriptome. A second pass with MAKER was run with the retrained SNAP, the trained AUGUSTUS and the similar set of input as above, and with these other options:

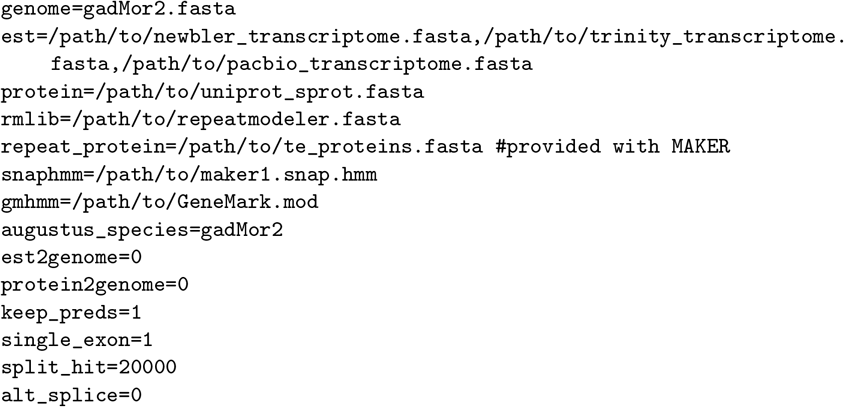

InterProScan version 5.4-47 [115] was run on the protein output of MAKER, providing gene ontologies and classifying protein domains and families. The protein output was BLASTed against SwissProt/UniProtKB release 2015_12, identifying putative gene names, with these options:

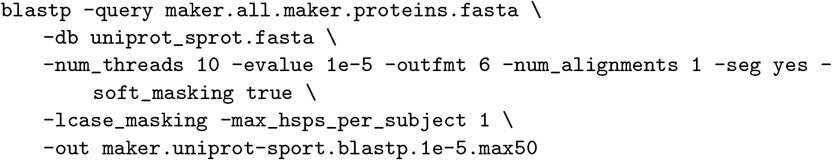

The final gene models set was filtered based on the AED score [116], including only gene models with an AED of less than 0.5. An AED of 0.0 would indicate that the gene model is in perfect accordance with its evidence (mRNA, EST, protein alignments) and 1.0 that it is not supported by any evidence.

### Investigating heterozygosity

To investigate the heterozygosity of this individual of Atlantic cod, we mapped the 300 bp insert size Illumina sequencing library to the genome assembly using bwa mem version 0.7.9a with the -M option [56]. Samtools version 1.1 was used to sort the bam files.

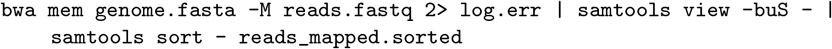

SNP and indel calling was done on the merged bam file using FreeBayes version v0.9.14-17-g7696787 [57], and SNP and indel calls with a quality >20 were kept with 'vcffilter -f “QUAL >20”'. Vcfstats was run on the resulting VCF file, giving the number of SNPs, MNPs, indels and complex regions.

We also mapped all PacBio reads using blasr from SMRT-Analysis 2.3.0, and called indels using PBHoney version r99 [59], annotating all indels larger than 20 bp. This numbered 70,278.

### Genome-wide short TR analysis

TRs of unit size 1–50 bp were detected with Phobos version 3.3.12 [18], options set were “-s 12 -outputFormat 0 -U 50”, i.e. requiring a minimum score of 12 for each TR, that is, the TR needed a score above 12, i.e. at least 13 mononucleotides, 7 dinucleotide, 5 trinucleotide repeat units, that is, minimum lengths of 13, 14 and 15 bp, respectively; Phobos native format as output; and up to a motif, or unit, size of 50 bp. A range of 1–50 bp was chosen in accordance with Mayer et al. [18]. A config file was then provided for the sat-stat version 1.3.12 program, yielding a diverse output of file with different statistics and a gff file:

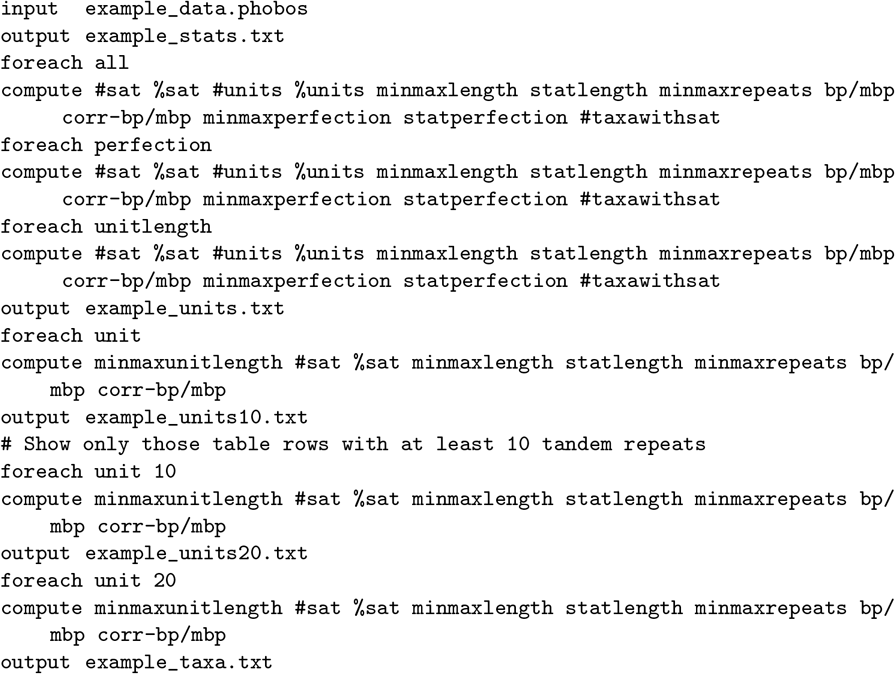

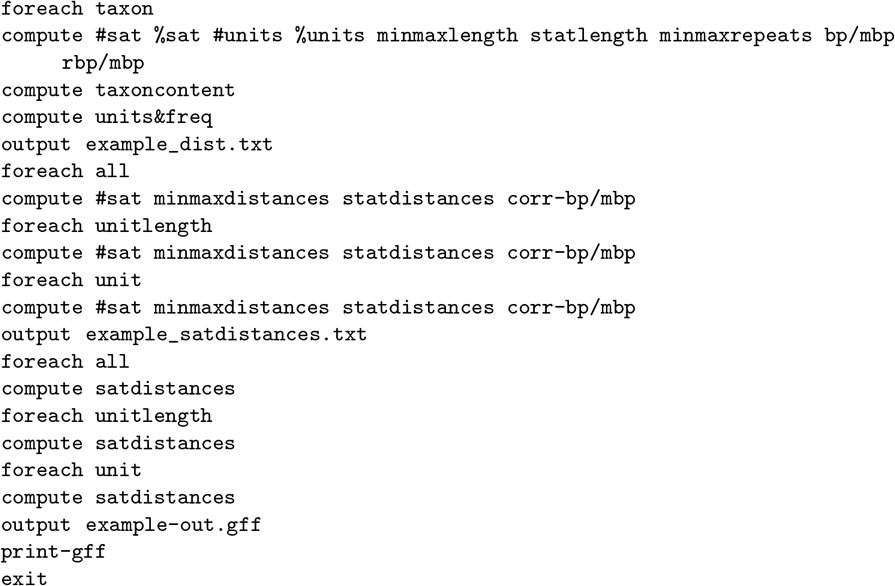

In addition, STRs were detected with lobSTR 4.0. First, TRF version 4.07b was run on the genome assembly with these options “gadMor2.fasta 2 7 7 80 10 24 6 -f -d -h”, and the resulting gadMor2.fasta.2.7.7.80.10.24.6.dat file was converted to bed format with convert_trf_bed_lobstr.py. A lobSTR index was created with the bed file and the genome, and allelotype classified the STRs using the Illumina 300PE library previously mapped with BWA, using these options:

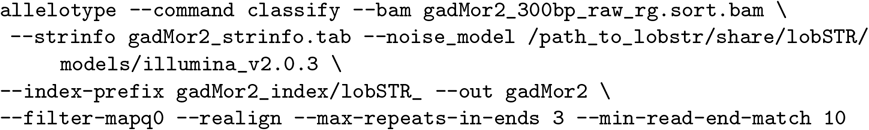

In addition to the different cod assemblies analyzed, we downloaded all assemblies from Ensembl release 81 (n = 68) (including Atlantic cod) and the California sea hare.

Star et al. [5] released three different assemblies, based on Newbler, Celera Assembler and a gene-model optimized, annotated version of the Newbler assembly, which is the one available from Ensembl and indicated herein as gadMor1. In gadMor1, contigs were reordered according to stickleback proteins during annotation, which resulted in significant improvements in regards to gene model construction compared with the original assembly. In all comparisons between different cod assemblies performed for this work, we compared to the gadMor1 assembly, since it is annotated and likely the one most used.

### Contig terminus analysis

Contigs from the assemblies of ALPILM, NEWB454, CA454PB, CA454ILM, gadMor1 and gadMor2 were created with the “cutN -n 1” command from seqtk version 1.0-r75, which cut at each gap (of at least one basepair, i.e. one or more Ns). The contigs were mapped against the gadMor2 assembly with BWA 0.7.12 and get_-positions_non_soft_hard_clip.py was used to create a BED file with only the edges of contigs that map uniquely with a mapping quality of 3 or more.

The intersect option from bedtools version 2.24.0 [117] was used to find overlaps between the contig termini and indels based on PBHoney tails output, SNPs, indels, MNPs and complex regions from mapping Illumina reads (300 bp insert size) to the genome, TRs called by Phobos, lack of coverage by Illumina, 454 and PacBio reads (zero depth as determined by mapped reads and bedtools genomecov), TEs and low complexity regions from RepeatMasker.

### Heterozygous TRs

We used bedtools [117] 2.24.0 to find the intersecting between the indels called by FreeBayes and PBHoney, and the TRs as annotated by Phobos. Indels were filtered based on depth (at least 5 reads) and genotype (0/1, heterozygous).

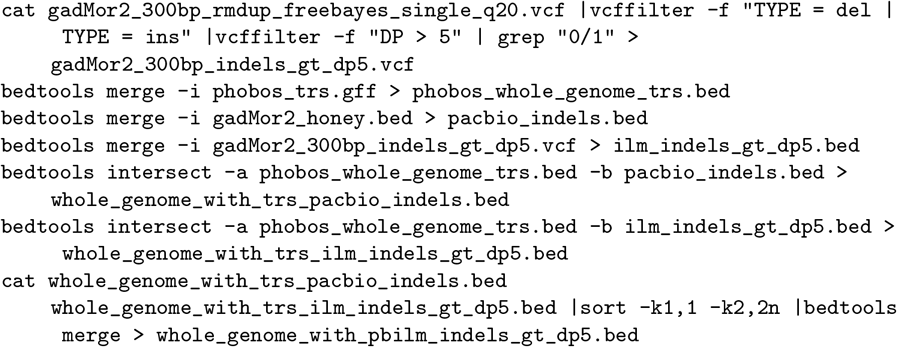

In the annotation of Atlantic cod, some genes were annotated that consist predominantly of TRs. Since these were in the annotation, they have some evidence in the form of protein or transcriptome alignment, and have an open reading frame. However, they seem to have no significant similarity with proteins from SwissProt/UniProtKB, and were removed based on this. This left 19,035 genes for this particular analysis.

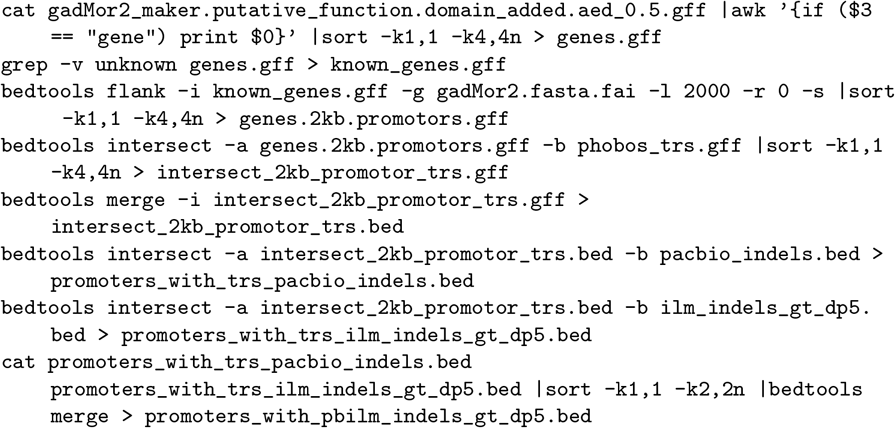

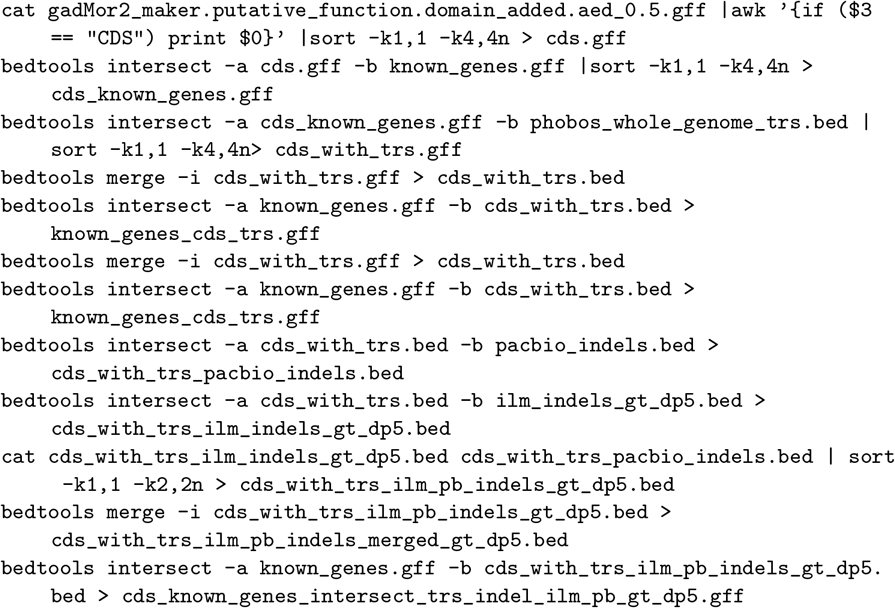

bedtools version 2.24.0 [117] was used to find the intersection between the coding sequence of genes with similarity to proteins from SwissProt/UniProtKB and TRs from Phobos. The result from this was intersected with indels called by FreeBayes (Illumina reads) and PBHoney (PacBio reads).

## Ethics approval and consent to participate

Not applicable.

## Consent for publication

Not applicable.

## Availability of data and material

Accession numbers at ENA (http://www.ebi.ac.uk/ena) for the gadMor2 assembly: WGS: CAEA020000001-CAEA020024862. Scaffolds: LN836027-LN845721. Chromosomes: LN845748-LN845770. Accession numbers for the Illumina libraries used for genome assembly: ERX1622640-ERX1622647. Accession numbers for the 454 libraries used for genome assembly: ERX1761596-ERX1761760. Accession numbers for the PacBio libraries used for genome assembly: ERX1787826-ERX1787972. Accession number for the PacBio Iso-Seq used for genome annotation: PRJEB18628.

The four original assemblies and gadMor2 available at Figshare: doi.org/10.6084/m9.figshare.3408247. All scripts used for analysis are available at https://github.com/uio-cels/cod2_scripts and https://github.com/uio-cels/Repeats.

## Authors contributions

OKT and AJN performed the genome assemblies. OKT and WBR annotated the genome assembly. OKT carried out the TR analysis. JRM, BPW and JK designed and wrote the assembly algorithms. JME, PP, AT-K and MS performed library creation and sequencing. HG and SL created the linkage map. RBE contributed samples for sequencing and evaluated assemblies. OKT, BS, SJ, BPW, KSJ and AJN wrote the paper with help from all the authors. All authors read and approved the final manuscript.

## Acknowledgements

All computational work was performed on the Abel Supercomputing Cluster (Norwegian metacenter for High Performance Computing (NOTUR) and the University of Oslo) operated by the Research Computing Services group at USIT, the University of Oslo IT-department (http://www.hpc.uio.no/). Sequencing library creation and high throughput sequencing was carried out at the Norwegian Sequencing Centre (NSC), University of Oslo, Norway; McGill University and Genome Quebec Innovation Centre, Canada and Pacific Biosciences of California Inc., Menlo Park. We are grateful for the willingness of Helle Tessand Baalsrud, Martin Malmstr0m and Monica H. Solbakken to investigate their favorite genes as a validation in different draft assemblies. Geir Kjetil Sandve provided essential help with implementing different algorithms for assembly reconciliation. We thank Jonas Korlach for his contribution to strategic aspects of the project and commenting on earlier versions of the manuscript, and Cassandra Trier for critical reading of the manuscript.

